# MCNV2 (Mendelian CNV Validation): Mendelian Precision for CNV quality assessment

**DOI:** 10.64898/2026.04.29.721462

**Authors:** Mame Seynabou Diop, Audrey Lemacon, Kuldeep Kumar, Benjamin Clark, Guillaume Huguet, Florian Bénitière, Martineau Jean-Louis, Sylvie Hamel, Sébastien Jacquemont

## Abstract

**Summary:** Detection of copy number variations from genomic sequencing and array data is prone to high false-positive rates. Distinguishing true variation from false positives remains challenging as quality metrics depend on technologies used, the quality of the data, and the calling algorithm. Mendelian inheritance in parent-offspring trios offers a powerful method to detect false positives, yet no tool exists to systematically compute, explore, optimize, and interpret the precision of CNV calls accordingly. Here we present Mendelian CNV Validation (MCNV2), an R package implementing Mendelian Precision (MP), as a reproducible metric for standardized CNV quality assessment. MCNV2 provides a command-line interface for pipeline integration and an interactive Shiny application for real-time exploration of MP across CNV types, size categories, and quality metrics.

**Availability and Implementation:** MCNV2 is available at https://github.com/JacquemontLab/MCNV2-Mendelian-CNV-Validation.

**Supplementary Information:** Supplementary information are available at https://mcnv2-mendelian-cnv-validation.readthedocs.io/en/latest/

## 1. Introduction

Copy number variations (CNVs) are a major source of genetic variation and contribute to many human diseases (Zarrei *et al*. 2015, Zou *et al*. 2026). Assessing the quality of CNV callset remains a key challenge, as no standardized metric exists to quantify the precision CNV calls across their characteristics and quality metrics (Lavrichenko *et al*. 2021). In addition, validating a novel CNV calling pipeline is also problematic as available ground truth datasets are small and incomplete (Yuan and Jia 2024).

Family-based validation using Mendelian inheritance in parent-offspring trios provides a robust method for CNV quality assessment. Indeed, most non-inherited CNVs in offspring represent false positives, as true *de novo* variants are rare (Porubsky *et al*. 2025). While several studies have leveraged this principle to validate their CNV calls (Yang *et al*. 2017, Lavrichenko *et al*. 2021), no standardized tool exists to systematically compute, explore, optimize, and interpret Mendelian Precision across CNV characteristics and quality metrics.

Here, we present Mendelian CNV Validation (MCNV2), an R package that formalizes Mendelian Precision as a reproducible metric to assess CNV quality. MCNV2 provides both a command-line interface for pipeline integration and an interactive Shiny application for real-time exploration of Mendelian Precision across CNV types, size categories, and quality filtering strategies. We demonstrate the utility of MCNV2 using CNV calls from WGS performed in 1 103 parent-offspring trios from the SPARK cohort (SPARK Consortium 2018). We show how CNV precision is impacted by quality metrics, gene coding and non-coding content as well as problematic genomic loci.

## 2. Methods

### 2.1 Overview of the MCNV2 framework

MCNV2 can be applied to CNVs from whole-genome sequencing (WGS), whole-exome sequencing (WES), or SNP array data. It requires CNVs in a standardized tabular format together with pedigree information of complete trios (both parents and an offspring).

The core principle of MCNV2 is Mendelian Precision (MP), defined as the proportion of inherited CNVs among all CNVs detected in an offspring. To compute MP, MCNV2 compares CNVs detected in offspring with those observed in their parents (Yang *et al*. 2017). CNVs shared with at least one parent are classified as inherited, whereas CNVs detected only in the offspring are classified as non-inherited:

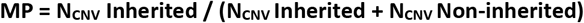

where **N**_**CNV**_ is the number of CNVs. Non-inherited includes true *de novo* variants and false positives. Given the low overall estimated rate (1.92%) of *de novo* CNVs in the general population (Zou *et al*. 2026), non-inherited CNVs in unconstrained genomic regions are predominantly false positives. MP therefore provides a reasonable estimate of call precision.

### 2.2 Workflow

The MCNV2 workflow consists of four steps (Fig 1A). First, CNVs are annotated with gene content from Gencode v45 (Frankish *et al*. 2021), gene-level constraint scores (LOEUF from gnomAD v4 (Gene constraint | gnomAD, n.d.), and problematic genomic regions from UCSC tracks (Kent *et al*. 2002), including segmental duplications, low-mappability regions, centromeres, and telomeres. Second, CNV inheritance status is based on reciprocal overlap between an individual and at least one parent. Overlap can be established at either the genomic coordinate level (CNV-level) or the gene-content level; the latter being more robust to breakpoint uncertainty due to large segmental duplications flanking many CNVs. Third, CNVs are filtered by size, quality metrics, concordance between CNV callers, and overlap with problematic regions identified in step 1. Fourth, MP is computed separately for deletions and duplications across seven CNV size categories and available quality metrics.

**Fig 1.**
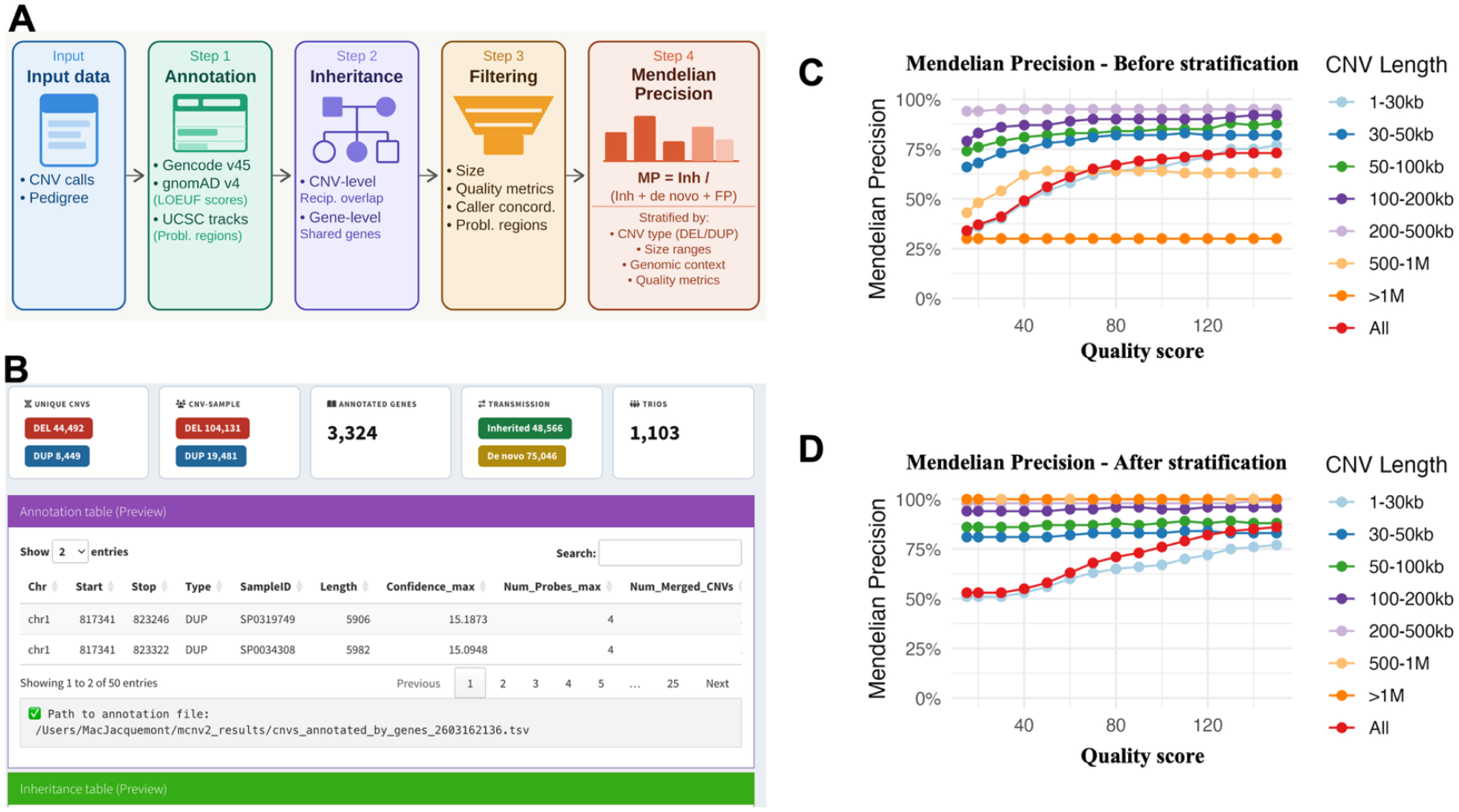
MCNV2 workflow and Mendelian Precision optimization. **(A)** MCNV2 four-step workflow: CNV calls and pedigree data are annotated with gene content (Gencode v45), LOEUF scores (gnomAD v4), and problematic genomic regions (UCSC tracks); inheritance is assigned by CNV type using either a CNV-level or gene-level approach; CNVs are filtered by size, quality metrics, caller concordance, and problematic region overlap; MP (Inherited / (Inherited + Non-inherited)) is computed by CNV type, size category, and quality metrics. **(B)** Shiny interface showing dataset summary statistics, annotated CNV table preview, and inheritance table preview. **(C)** MP for deletions versus quality score threshold, stratified by CNV size, without additional filtering. **(D)** MP for deletions after combined filters: exclusion of problematic regions, LOEUF stratification (LOEUF < 0.6), and algorithm concordance (PennCNV and QuantiSNP).

### 2.3 Identifying CNVs at high probability of occurring *de novo*

Non-inherited CNVs include both false positives and true *de novo* events. Since the probability of occurring *de novo* is correlated with the number of genes under selective constraint within a CNV (Huguet *et al*. 2018), such CNVs disrupting one or more constrained genes systematically reduce MP despite being true positives.

MCNV2 therefore annotates each CNV with LOEUF (Loss-of-function Observed/Expected Upper bound Fraction), a gene-level measure of intolerance to loss-of-function mutations from gnomAD v4 (Gene constraint | gnomAD, n.d.), and excludes CNVs overlapping at least one constrained gene from MP computation (LOEUF below a user-defined threshold).

### 2.4 Interfaces

MCNV2 is provided through two complementary interfaces: a command-line interface for automated pipeline integration and large-scale analyses, and an interactive Shiny application for real-time exploration and visualization of MP across filtering strategies.

### 2.5 Computational performance and resource usage

Processing 1 103 trios (123 612 offspring CNVs) was completed in under 120 seconds, with peak memory usage below 8 GB on a MacBook Pro (2019, macOS Sonoma 14.7.6, 2.3 GHz 8-Core Intel Core i9, 64 GB RAM) running R version 4.5.1.

## 3. Results

### 3.1 Application of MCNV2 to the SPARK WGS cohort

We applied MCNV2 to 1 103 parent–offspring trios from the SPARK whole-genome sequencing cohort (~30× coverage) (SPARK Consortium 2018). CNVs were detected using PennCNV (Wang *et al*. 2007) and QuantiSNP (Colella *et al*. 2007) through two pipelines (Bénitière *et al*. 2026, Diop 2026) and yielded 123 612 CNVs (quality score ≥15) in offspring.

A summary of dataset statistics and a preview of the annotated CNV and inheritance tables are provided (Fig. 1B). Results are shown for deletions (Fig 1C and 1D).

### 3.2 Increasing CNV quality score improves Mendelian Precision

Inheritance status was assigned using a CNV-level approach with 50% reciprocal overlap, and MP was computed across seven CNV size categories (Fig 1C). MP increased with higher CNV quality score thresholds across all size categories, as expected given that higher quality scores reflect greater confidence in true CNV calls (Macé *et al*. 2016). Larger deletions (50-500 kb) showed consistently high MP, improving from ~74% to ~92%. Smaller deletions showed the greatest improvement; 1-30 kb deletions improved from ~33% to ~77%, and 30-50 kb deletions from ~66% to ~82%. MP reaches a plateau beyond a certain quality score threshold, beyond which stricter filtering reduces the number of CNVs without further improving precision.

### 3.3 Improving Mendelian Precision through artifact removal and *de novo* stratification

CNV calling is unreliable in problematic regions (as defined by the UCSC Genome Browser) (Kent *et al*. 2002). Removing CNVs in the latter regions is essential before MP assessment.

To distinguish false positives from true *de novo*, CNVs overlapping constrained genes (LOEUF < 0.6, as defined by (Gene constraint | gnomAD, n.d.)) are excluded from MP computation. This improves MP from ~27-60% to 100% for large deletions > 500kb (Fig 1D).

Calling algorithms have different statistical models, making false positives typically algorithm specific, while true CNVs are more likely to be called by both algorithms. Algorithm concordance, requiring CNVs to be called by at least two callers with a reciprocal overlap of at least 50%, improved MP across all size categories compared to unfiltered results (Fig. 1C). Larger deletions (50–500 kb) improved from ~74-92% to ~82-94%, smaller deletions (1-30 kb) from ~33-77% to ~51-69%, and 30-50 kb deletions from ~66-82% to ~79-83%.

After applying the steps described above, MP improved substantially across all size categories and continues to improve with higher quality score values (Fig 1D). We also demonstrate that MP varies across size categories, with smaller CNVs (< 30kb) requiring higher quality scores to achieve MPs similar to larger CNVs. Comparison with the unfiltered results (Fig. 1C) illustrates the cumulative impact of these filters on call precision across all size categories.

### 3.4 Best-practice filtering strategies

The highest MP in an artifact-free dataset would be the 1-frequency of *de novo* events, approximately 98%, given the frequency of *de novo* CNVs is, on average, 1.92% (Zou *et al*. 2026) in the general population and varies across CNV sizes and gene content. Datasets with MPs exceeding 70% are recommended for reliable downstream analyses.

We recommend applying filters sequentially to assess the individual contribution of each filter to MP: first, excluding CNVs overlapping problematic genomic regions, then applying LOEUF-based stratification to separate *de novo* events from false positives, then filtering by CNV size and quality score thresholds. Algorithm concordance, when available, provides the largest single improvement in MP.

### 3.5 CNV-level versus gene-level analysis

MCNV2 supports both CNV-level (coordinate-based) and gene-level (gene-based) inheritance definitions. In this Application Note, we focused on CNV-level analysis because we used whole-genome sequencing, allowing coding and non-coding CNV identification. In the context of CNVs called using technologies such as exome, gene-level MP can be used. Detailed comparisons are provided in the online documentation.

## 4. Conclusion

MCNV2 provides a framework for CNV quality assessment in parent-offspring trio datasets using Mendelian inheritance and a prior knowledge of the frequency of *de novo* CNVs. It presents MP values across CNV quality metrics and size categories, helping researchers select thresholds providing the best trade-off between precision and number of CNVs for their downstream analyses. Beyond quality control, MCNV2 serves as a benchmark-free validation framework for CNV detection algorithms.

## Acknowledgements

This research was enabled by support provided by the Digital Research Alliance of Canada (https://www.alliancecan.ca/). We are grateful to all of the families in SPARK, the SPARK clinical sites, and SPARK staff. We appreciate obtaining access to phenotypic and genetic data on SFARI Base. Approved researchers can obtain the SPARK population dataset described in this study by applying at https://base.sfari.org. We thank Thomas Renne for his time and constructive feedback during the evaluation of the application.

